# Emerin modulation impacts viability, proliferation, migration, and DNA repair signaling in cisplatin-treated glioblastoma cells

**DOI:** 10.64898/2026.06.25.734655

**Authors:** Donna Joe Farfan Hilares, Fábio Luís Forti

## Abstract

Emerin (EMD), an inner nuclear membrane protein essential for nuclear architecture integrity, gene expression, cellular signaling, and chromatin stability, interacts with the LINC complex and participates in cytoskeleton-nucleoskeleton communication by binding to nuclear actin filaments. EMD is implicated in migration, invasion, and metastasis in some tumors, but its role in glioblastoma (GBM) remains unclear. This study evaluated the effects of EMD knockdown and overexpression in GBM cell lines following genotoxic treatment with cisplatin. In both wild-type p53 (U87-MG) and mutant p53 (U138-MG) GBM cells, EMD expression is high, and cisplatin treatment did not affect these protein levels. EMD knockdown in U87-MG cells significantly increased cisplatin IC50, viability, and proliferation. Conversely, stable overexpression of EMD in U87-MG cells led to reduced cisplatin IC50, viability, proliferation, and migration. EMD knockdown or overexpression did not affect any U138-MG phenotypes, with or without cisplatin treatment. Modulation of EMD levels causes morphological changes in stress fiber cytoskeleton, whereas overexpression of EMD in U87-MG cells promotes an increase and a decrease in nuclear and cytoplasmic actin levels, respectively. These biological responses of U87-MG cells overexpressing EMD were coincidentally associated with alterations in the levels of pH2AX(Ser139), p-p53(Ser15), p53, and p21Kip1 proteins after cisplatin exposure. In sum, modulation of EMD levels affects the viability, migration, and proliferation of wild-type p53 GBM cells treated with cisplatin, suggesting unknown roles in the DNA damage response and repair. This work highlights EMD as a potential regulator of GBM chemoresistance and a target for therapeutic intervention.

## Introduction

Emerin is a ubiquitously expressed protein localized at the inner nuclear membrane, associated with several important nuclear processes, including regulation of gene expression, cellular signaling, chromatin condensation and organization, and nuclear architecture [1][2]. For many years, its study has been primarily focused on the context of nuclear envelopathies, as mutations in the EMD gene are the main cause of Emery–Dreifuss muscular dystrophy (EDMD), a disorder characterized by defects in skeletal muscle, cardiac conduction abnormalities, and premature aging [3][4]. However, over the past 15 years, Emerin has gained increasing attention in the oncological context [5][6][7][8][9], particularly as a potential mediator of nuclear shape stability in prostate, lung, and breast cancers, and has even been proposed as an effective clinical diagnostic marker [10].

In this context, Emerin is not only relevant for its role in maintaining nuclear architecture. Within the nucleoplasm, it interacts with nuclear actin, binding both globular [11] and filamentous actin [12]. In vitro studies have demonstrated that Emerin caps the pointed end of F-actin filaments, stabilizing them and promoting their polymerization [12]. Thus, it may contribute to the formation of a nuclear actin network that supports nuclear architecture integrity and chromatin organization, processes that are particularly relevant for the DNA damage response [13]. At the inner nuclear membrane, Emerin also interacts with SUN1 and SUN2, SUN domain-containing proteins (Sad-1 and UNC-84) that are components of the LINC (Linker of Nucleoskeleton and Cytoskeleton) complex, which physically connects the nucleoskeleton to the cytoskeleton [14][15][16]. Through these interactions, Emerin may act as a bridge between cytoplasmic and nuclear actin.

Recent studies have demonstrated that modulation of cytoskeleton-associated pathways, such as Rho GTPase signaling and its downstream effectors ROCK kinases, directly influences cell survival and DNA repair mechanisms in a p53-dependent manner [17]. In this context, ROCK inhibition has been shown to exert opposite effects in glioblastoma (GBM) cells, reducing DNA repair capacity and survival in p53 wild-type cells, while enhancing repair efficiency and resistance in p53-mutant cells through differential modulation of pathways such as non-homologous end joining (NHEJ), nucleotide excision repair (NER), and homologous recombination [17]. Furthermore, recent evidence highlights that actin cytoskeleton dynamics, regulated by Rho GTPases, play a central role in both the development and reversal of chemoresistance in GBM [18]. Temozolomide- and cisplatin-resistant cells exhibit increased Rho pathway activity, associated with enhanced double-strand break repair and reprogramming of DNA repair pathways, including reduced levels of mismatch repair proteins. Notably, inhibition of this pathway restores sensitivity to chemotherapeutic agents, reversing the resistant phenotype in a p53-dependent manner [18]. Together, these findings indicate that cytoskeletal dynamics and nucleus–cytoskeleton communication constitute a critical axis in the regulation of DNA damage response and therapeutic resistance in GBM.

Despite its emerging relevance in oncology, the role of Emerin in glioblastoma (GBM) remains poorly explored, particularly in the context of genotoxic stress. GBM accounts for 48.6% of malignant tumors of the central nervous system [19] and is characterized by highly aggressive biology, with elevated proliferative and infiltrative capacity, as well as significant genetic and molecular heterogeneity, contributing to recurrence, therapy resistance, and rapid progression [20]. As a result, prognosis remains poor, with a median overall survival of less than 15 months [21] and 5-year survival rates below 10% [20]. In this context, identifying novel molecular mechanisms involved in tumor response to genotoxic stress is essential for the development of more effective therapeutic strategies. Therefore, in the present study, we investigated the effects of Emerin knockdown and overexpression in glioblastoma cell lines with different p53 statuses following cisplatin treatment.

## Materials and Methods

### Bioinformatic analysis

Genomic alterations and gene expression analyses of *EMERIN*, LINC complex, and nuclear lamina genes were performed using publicly available datasets. Genomic alteration profiles were obtained from the Glioblastoma Multiforme cohort (TCGA, PanCancer Atlas) through the cBioPortal platform. Gene expression analyses were conducted using GEPIA, based on data from The Cancer Genome Atlas (TCGA) PanCancer Atlas for glioblastoma (GBM) and low-grade glioma (LGG), complemented with normal brain tissue data from the Genotype-Tissue Expression (GTEx) project.

### Cell cultures

U87-MG (ATCC # HTB-14) is a cell line with epithelial morphology isolated from malignant gliomas from the brain of a human male patient expressing wild-type tp53 gene. U138-MG (ATCC# HTB-16) is a polygonal cell line isolated from the brain of human male patient with glioblastoma, a glioma of histological grade IV, which expresses the tp53 gene harboring the R175H mutation. Both cell lines were obtained from ATCC in 2020 by Prof. Dr. Carlos Frederico Martins Menck (ICB-USP) and Prof. Dr. Elza Tiemi Sakamoto Hojo (FFCLRP-USP) and kindly donated to our laboratory. Cells were maintained in DMEM culture medium (Invitrogen) supplemented with 10% fetal bovine serum (Cultilab), ampicillin (25 µg/mL), and streptomycin (100 µg/mL), incubated at 37°C in a humidified atmosphere containing 5% COL (according to the handling information provided by the ATCC repository), and maintained in culture only up to 20^th^ passage. The authentication of our cell lines is done annually using the STR (Short Tandem Repeat) methodology, a molecular biology technique based on the identification of polymorphic regions present in the sample’s genome. This technique was developed for forensic applications and is used by international collections such as the American Type Culture Collection (ATCC), among others, and we consider a match of at least 90% between the STR profile of the GBM cells in relation to a reference sample. Every six months all cell lines of our laboratory are tested for mycoplasma contamination using the Myco-sniff^TM^ mycoplasma PCR detection kit and treated for 15 days with mycoplasma removal agent Plasmocin® according to the manufacturer protocol (InvivoGen).

### Cisplatin treatments

Cisplatin (Sigma-Aldrich) was dissolved in 0.9% NaCl saline solution at a concentration of 10 mM and stored at 4°C for up to 3 months. The drug was further diluted in DMEM immediately prior to application to the cells.

### siRNA transfection and gene knockdown

Emerin gene knockdown was performed by transient transfection of Emerin-specific small interfering RNA (MISSION® siRNA; Sigma-Aldrich) using Lipofectamine RNAiMAX (Invitrogen) as the transfection reagent. Knockdown efficiency was validated by Western blot analysis and the maximum knockdown reached by 72 hours after transfection. Cisplatin treatment was carried out 48–72 hours post-transfection.

### Generation and selection of stable eGFP and eGFP-EMD–expressing cell lines

To generate U87-MG and U138-MG cell populations stably expressing eGFP (control) or eGFP-Emerin (GFP-EMD), the eukaryotic expression vectors pEGFP-C1 and pEGFP-C1-EMD (previously clones in our laboratory, unpublished data) were transfected using Lipofectamine 3000 with P3000 reagent (Invitrogen). Seventy-two hours post-transfection, selection was initiated by culturing the cells in a medium supplemented with geneticin (G418; Gibco) at a concentration of 1 mg/mL. The selective medium was replaced every three days to maintain selection pressure and eliminate non-transfected cells. Selection efficiency and eGFP expression were monitored using an Olympus IX51 inverted fluorescence microscope controlled by CellSens software (Olympus). Stable expression of Emerin was further validated by Western blot analysis. After selection and characterization, clonal cell lines were maintained in culture medium supplemented with geneticin (250 µg/mL).

### Western Blot Analysis

Cell lysates were prepared using RIPA buffer (50 mM Tris-HCl pH 7.2, 1% Triton X-100, 0.5% sodium deoxycholate, 0.1% SDS, 500 mM NaCl, 10 mM MgClL) supplemented with protease inhibitors (2 µg/mL leupeptin, 2 µg/mL pepstatin A, 2 µg/mL aprotinin, and 1 mM PMSF) and phosphatase inhibitors (1 mM NaLVOL and 1 mM NaF), followed by centrifugation (15 min, 13,500 rpm, 4°C). Protein concentration was determined at 595nm using the Bradford assay in a Varioskan microplate reader (ThermoFisher Scientific). Proteins were separated by SDS-PAGE and transferred to nitrocellulose membranes (Millipore) using a wet transfer system (Mini Trans-Blot, Bio-Rad). Membranes were stained with Ponceau S, blocked with 5% non-fat milk in TBS-T, and incubated with the appropriate primary and secondary antibodies. Detection was performed using the Odyssey (LI-COR) system, and densitometric analysis was carried out with Empiria Studio software (LI-COR).

### MTT cell viability assay

Cell viability under the experimental conditions was assessed using the MTT reduction assay [3-(4,5-dimethylthiazol-2-yl)-2,5-diphenyl tetrazolium bromide]. In this assay, cells were treated with 2.5, 5, 10, 20, and 25 µM cisplatin for 48 hours. After treatment, MTT (Invitrogen) was added at a final concentration of 500 µg/mL and incubated for 3 hours at 37°C. The resulting formazan crystals were solubilized in DMSO, and absorbance was measured at 570 nm in a Varioskan microplate reader (ThermoFisher Scientific). Dose–response curves were generated, and ICLL values were calculated using GraphPad Prism 8.0 software.

### Clonogenic survival assay

For the analysis of cell survival, cells were seeded in 35-mm plates and treated with cisplatin at 2 µM (U87-MG) or 1 µM (U138-MG) for 24 hours. Colony formation was monitored for 2 weeks, with culture medium replaced every 2 days. Cells were fixed with 10% formaldehyde in PBS for 10 minutes and stained with 0.01% crystal violet for 10 minutes. For quantification, the stained cells were solubilized with 33% acetic acid, and absorbance was measured at 590 nm in a Varioskan microplate reader (ThermoFisher Scientific).

### 3D spheroid culture

For tumorsphere formation, 96-well plates were pre-coated with 1.5% agarose prepared in DMEM. Subsequently, 2 × 10L cells were seeded per well and cultured for 72 hours. DMEM supplemented with 10% FBS was centrifuged for 10 minutes prior to use to ensure the removal of debris and unwanted aggregates that could interfere with spheroid formation.

### 2D radial spheroid migration assay

To assess cell motility capacity in 2D substrates, tumorspheres were transferred to 96-well tissue culture–treated plates to allow cell adhesion and radial outgrowth and subsequently treated with 50 µM cisplatin. Cell migration was monitored for 48 hours. The migration area was quantified using FIJI software (ImageJ).

### Statistical analysis

At least three independent experiments were performed, each with two to four technical replicates. Statistical analyses were conducted using two-way analysis of variance (two-way ANOVA) in GraphPad Prism version 11.0 (GraphPad Software, San Diego, CA, USA). Asterisks indicate statistical significance (*p < 0.05, **p < 0.01, ***p < 0.001, and **p < 0.0001).

## Results

### Bioinformatics datamining of *EMERIN* and LINC genes in gliomas

The linker of nucleoskeleton and cytoskeleton (LINC) is a protein complex associated with both inner and outer membranes of the nucleus. Major protein components surrounding LINC are Lamins A-C and Emerin, which indirectly connect the structural cytoskeleton with chromatin. A set of ten genes related to these proteins was investigated as to their genetic frequency in samples of gliomas from patients: no alteration was predominantly found, while some genes presented some gain of function (*LAMIN B2, SUN1* and *NESPRIN 4*) or shallow deletion (*SUN2, NESPRIN 1, 2 , 3,* and *EMERIN*) (Fig. 1A). Some of these genes present differential gene expressions (DGE) when comparing low grade gliomas (LGG) with high grade gliomas (GBM); for example, all three *LAMIN* genes are upregulated in GBM, while *NESPRIN 1* is the most downregulated gene (Fig. 1B); all other genes are subtly differentially expressed in both gliomas. Similarly, in terms of overall survival (OS) of patients, high expression of *EMERIN* (Fig. 1C), *LAMIN A, LAMIN B1, LAMIN B2, SUN1*, and *NESPRIN 4* correlates with lower OS, low expression of *NESPRIN 1* and *NESPRIN 2* is correlated with higher OS (Fig. S1), whereas *SUN2* and *NESPRIN 3* changes the patient OS along the time (Fig. 1C and Fig. S1). In addition to this set of genes, we included *TP53* and *TP53I13* and checked for their co-expression correlation with the *EMERIN* gene (Fig. S2). Both Spearman and Pearson coefficients showed a positive coefficient for the pairs *EMERIN* and *LAMIN A*, *TP53* and *TP53I13*, but no correlation was found for all other genes, even negative ones.

**Figure 1.**
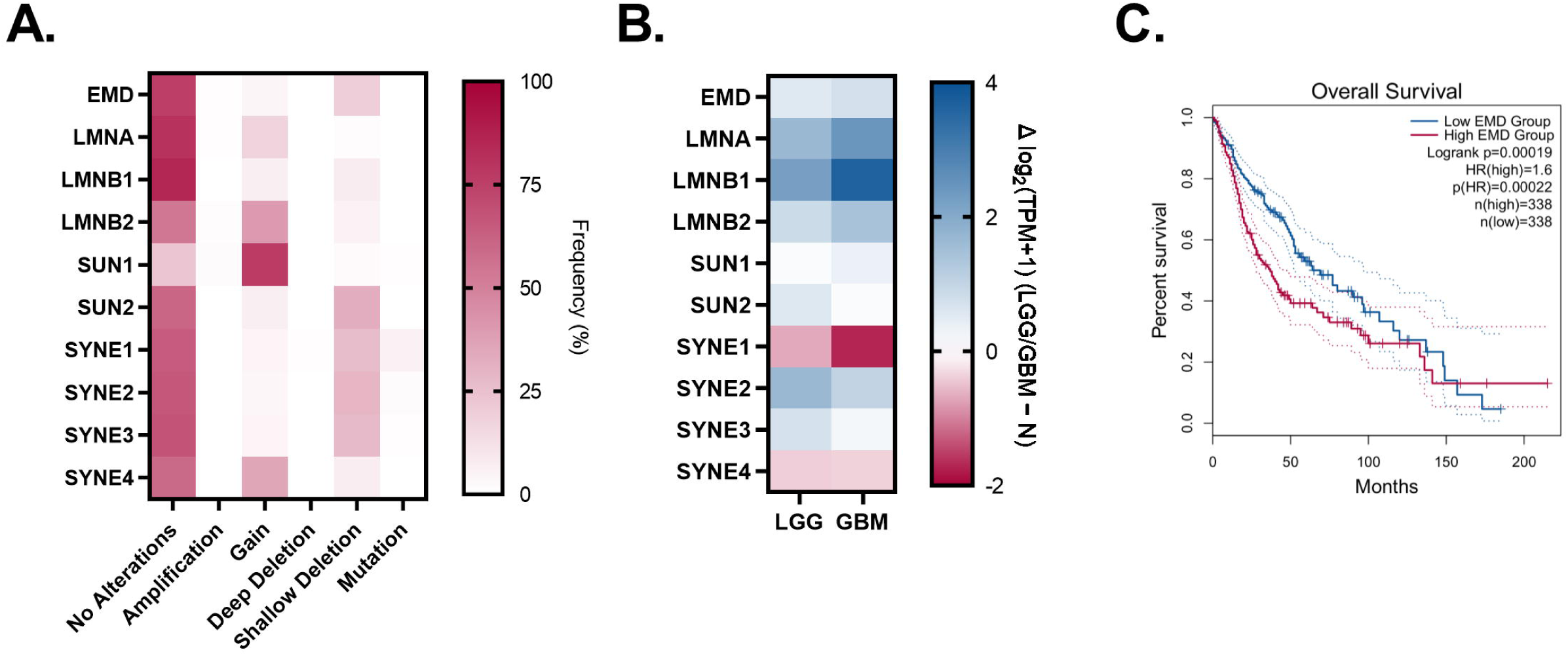
Genomic alterations, gene expression profiles, and overall survival analysis of Emerin, LINC complex, and nuclear lamina components in glioma (LGG and GBM). **A**. Heatmap showing the frequency of genomic alterations (amplification, gain, deep deletion, shallow deletion, and mutation). Darker shades indicate a higher frequency of alterations. B. Heatmap representing relative mRNA expression levels in low-grade glioma (LGG) and glioblastoma (GBM) compared to normal brain tissue (N). The color scale indicates logL(TPM+1) values calculated as LGG–N and GBM–N. Darker shades represent higher relative expression compared to normal tissue. C. Kaplan–Meier overall survival (OS) analysis of GBM and LGG patients stratified according to high (red) and low (blue) gene expression levels of EMD. Hazard ratio (HR), log-rank p-value, and number of patients in each group (n) are indicated within each plot. Dashed lines represent 95% confidence intervals. Genes abbreviation: *EMD: EMERIN; LMNA: LAMIN A/C LMNB1-2: LAMIN B1-B2: SYNE1-4: NESPRIN 1-4*.

### Modulating Emerin expression

According to the previous data and considering its unknown involvement on DNA repair of cells under genotoxic stress, we decided to modulate the expression of Emerin protein in glioblastoma cells. Two cell lines with different background of the tp53 gene were selected, U87-MG (wt-p53) and U138-MG (mut-p53), and submitted to siRNA transient transfection or exogenous expression of eGFP-Emerin (eGFP-EMD) construct (Fig. 2). The knockdown of both cells with siEMD was successfully achieved (Fig. 2A and 2B, and Fig. S2A and S2B) compared to parental cells or control siRNA. Similarly, it succeeded the ectopic overexpression of Emerin (Fig. 2C and 2D, and Fig. S2C and S2D) with eGFP-EMD, which can be preferentially located at the nuclear membrane, as expected and shown by immunofluorescence staining (Fig. 2E).

**Figure 2.**
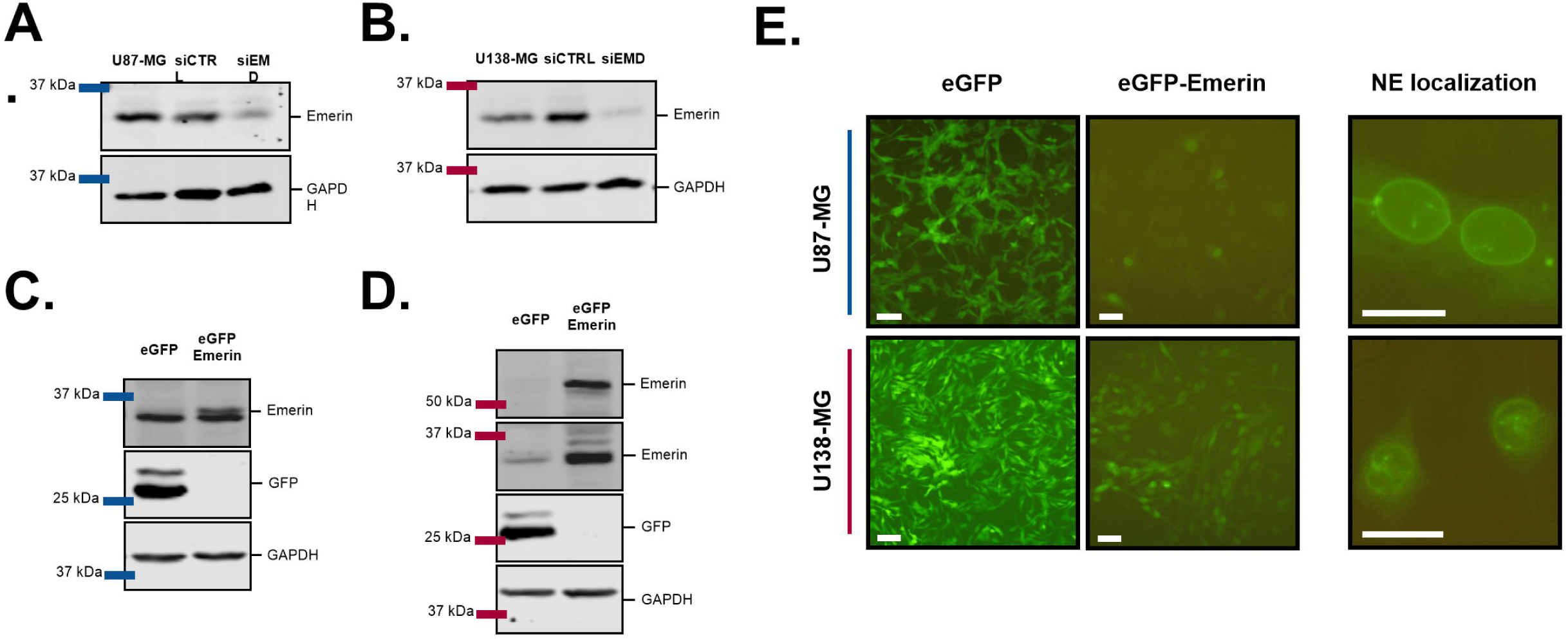
Validation of Emerin knockdown and ectopic overexpression in glioblastoma cell lines. A–B. Validation of Emerin knockdown by siRNA in U87-MG (A) and U138-MG (B) cells. Cells were transfected with control siRNA (siCTRL) or Emerin-specific siRNA (siEMD), and protein levels were analyzed by Western blot. **C–D.** Validation of stable expression of eGFP (control) or eGFP-Emerin in U87-MG (C) and U138-MG (D) cells by Western blot analysis. GAPDH was used as a loading control in all immunoblots. **E.** Representative fluorescence microscopy images of U87-MG and U138-MG cells expressing eGFP or eGFP-Emerin. The right panels highlight nuclear envelope (NE) localization of eGFP-Emerin. Representative micrographs were obtained using 20x and 40x objective lenses. Scale bars represent 100 µm for 20x images and 25 µm for 40x images.

### Down and upregulation of Emerin protects and sensitizes wt-p53 GBM, respectively, after genotoxic stress by cisplatin

Taking advantage of successful modulation of Emerin expression in both GBM cell lines, we next treated them with chemotherapeutic genotoxin cisplatin (Fig. 3). The previous silencing of U87-MG cells by siEMD and subsequent treatment with cisplatin protect cells from the damage by increasing viability (up to 25µM of cisplatin) and by promoting survival (at 2µM of cisplatin) in comparison to the parental cells (Fig. 3A, 3E and 3I). On the other hand, cells stably overexpressing eGFP-EMD were sensitive to up to 25µM of cisplatin and unable to recover survival after 2µM of cisplatin treatments (Fig. 3B, 3F and 3J, and Fig. S3A and S3C). Similar assays of viability and proliferation were performed with U138-MG cells; however, and interestingly, either silencing or overexpressing Emerin did not change the cells response to cisplatin (Fig. 3C, 3D, 3G, 3H, 3K and 3L, and Fig. S3B and S3D).

**Figure 3.**
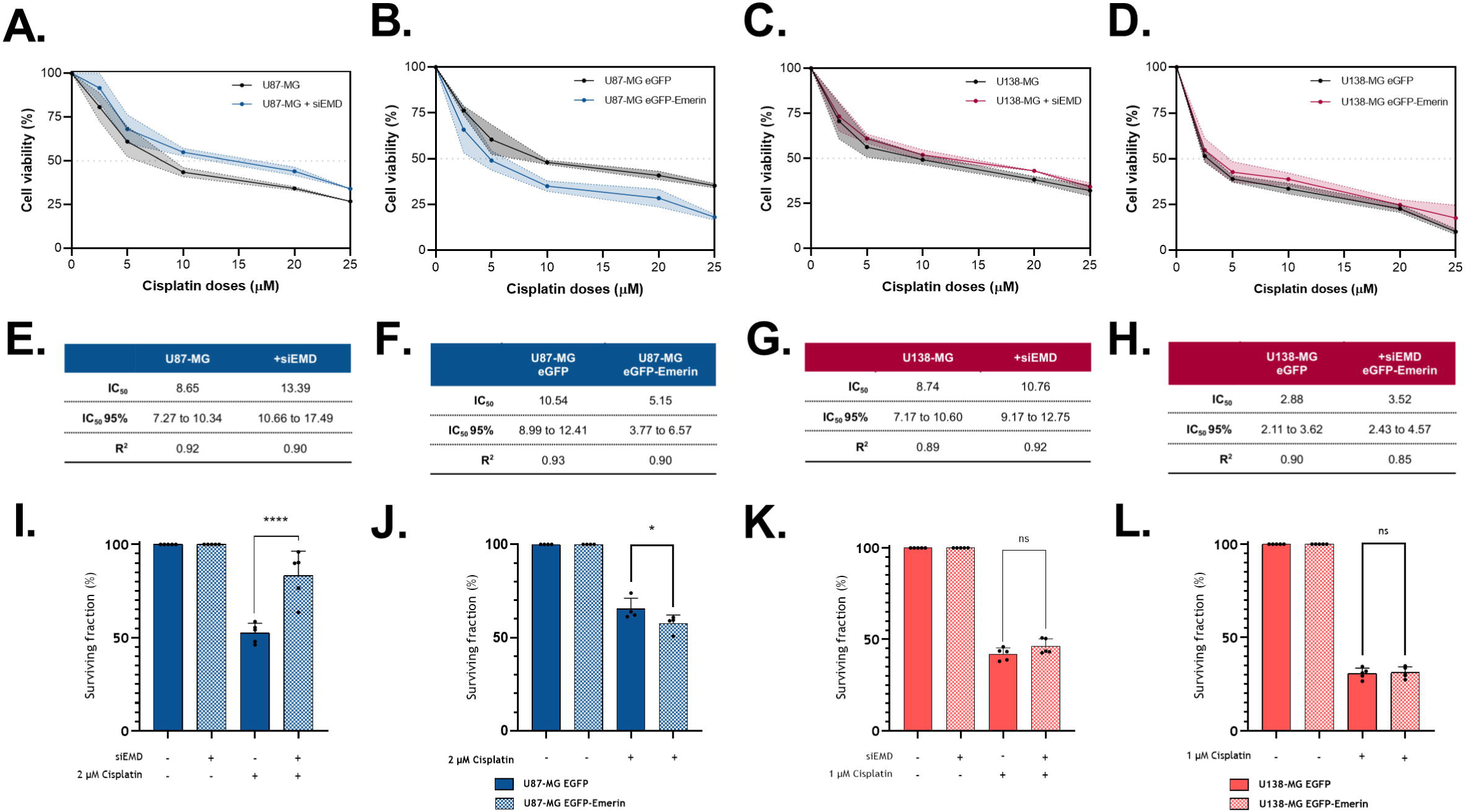
Emerin modulation affects viability and survival in p53 wild-type GBM cells following cisplatin treatment. A–D. MTT cell viability assay in U87-MG (A–B) and U138-MG (C–D) cells following Emerin modulation and cisplatin treatment. **E–H.** Tables showing IC□□ values for cisplatin, R² coefficients, and 95% confidence intervals for U87-MG (E–F) and U138-MG (G–H) cells. **(I–L)** Clonogenic survival fraction of U87-MG (I–J) and U138-MG (K–L) cells after Emerin modulation and cisplatin treatment. Graphs show SD from three independent experiments, with individual replicate values displayed. Statistical analysis was performed using two-way ANOVA.

### Emerin overexpression changes the GBM motility in spheroid migration assays

Since wt-p53 GBM cells overexpressing Emerin are more sensitive to cisplatin treatments, we decided to check whether migration of GBM cells is also affected by Emerin levels. Both U87-MG and U138-MG cells overexpressing or not eGFP-EMD were grown in agarose-covered 96 wells plates and cultured for 3 consecutive days until formation of physiologically relevant 3D spheroids (≥300nm). After that, spheroids were transferred to cell culture plates, treated by 50µM cisplatin, and followed by 24h and 48h after. Interestingly, ectopic overexpression of Emerin reduces the migration capacity of U87-MG cells especially 48h after (Fig. 4A and 4C), while in U138-MG cells it causes an increase in their migration in 48h after (Fig. 4B and 4C). These results are independent of the cisplatin treatments, which always cause similar reduced invasion of both GBMs compared to untreated cells very likely by reducing cell proliferation.

**Figure 4.**
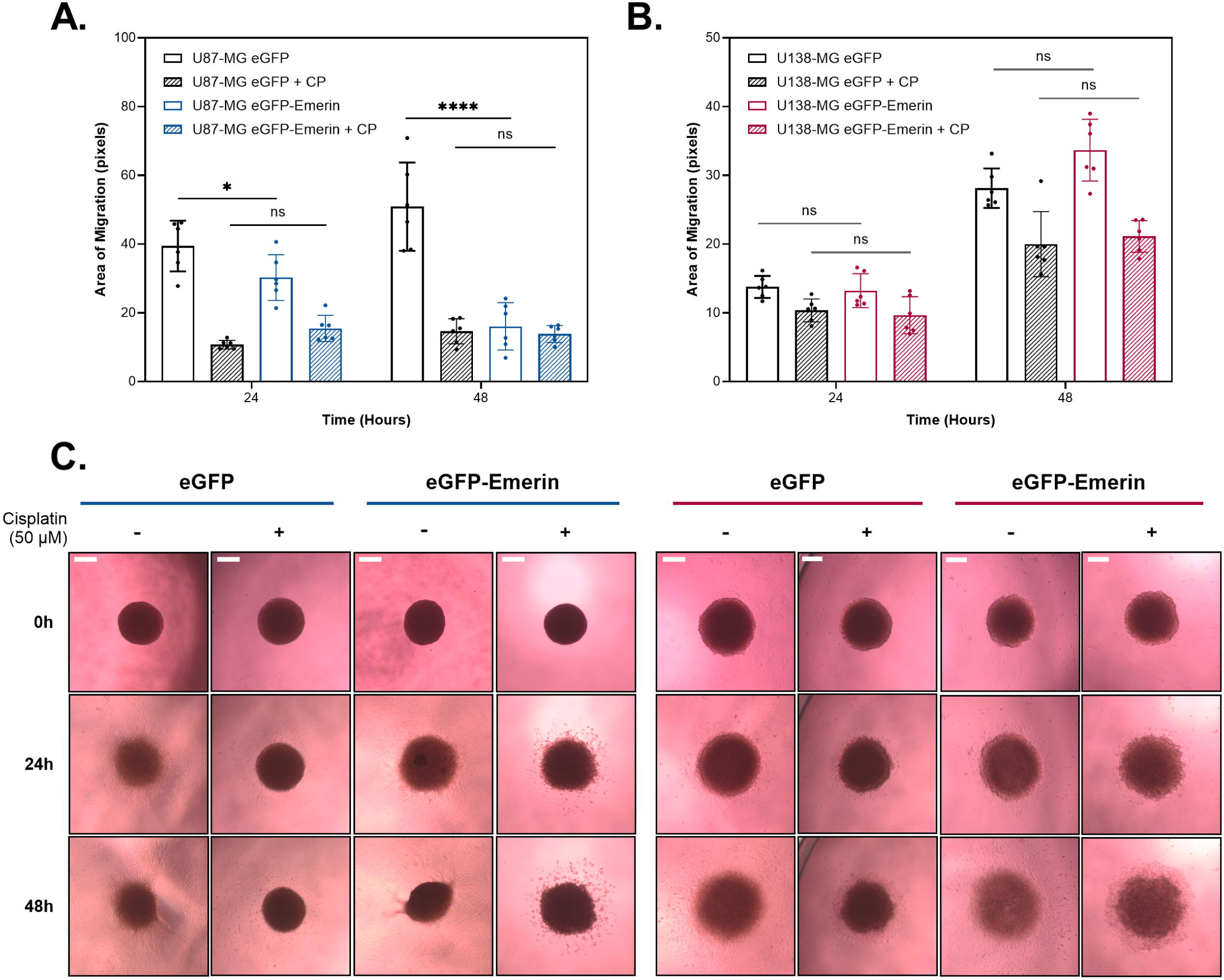
Emerin overexpression impairs migration in p53 wild-type GBM cells following cisplatin treatment. **A–B.** Quantification of radial migration area from tumorspheres of U87-MG (A) and U138-MG (B) cells stably expressing eGFP (control) or eGFP-Emerin, treated or not with 50 µM cisplatin. Migration was measured at 24 and 48 hours relative to zero-hour timepoint and expressed as invaded radial area (pixels). Data is presented as mean ± SD with individual data points from independent experiments. Statistical analysis was performed using two-way ANOVA. Significance is indicated as *p < 0.05; ***p < 0.0001; ns, not significant. **C.** Representative bright-field images of tumorspheres from U87-MG and U138-MG cells expressing eGFP or eGFP-Emerin, treated or not with 50 µM cisplatin by 0, 24, and 48 hours. Representative micrographs were obtained upon magnification of 4x objective lenses. Scale bar = 200 µm.

### Nucleocytoplasmic actin levels are altered by modulation of Emerin expression

To assess whether different status of p53 could affect cytoplasmic and nuclear actin distribution in both GBM cell lines, as well the ectopic Emerin overexpression, cell fractionation experiments were performed followed by immunoblottings (Fig. 5). Constitutive expressions of vinculin and NAT10 proteins were assumed as positive cytoplasmic and nuclear controls, respectively. By overexpressing Emerin, U87-MG cells presented reduced levels of cytoplasmic actin (Fig. 5A and 5E) and increased levels of nuclear actin (Fig. 5B and 5E), whilst U138-MG cells tend to show the opposite (Fig. 5C, 5D and 5F), just like expected for wild-type and mutated p53 cells, respectively. Additionally, imbalance of Emerin levels in both cell lines, either by overexpressing or knocking down, promotes visible changes in size and distribution of actin stress fibers, with augmented fragmentation and/or aggregation in comparison to control cells (Fig. 5G).

**Figure 5.**
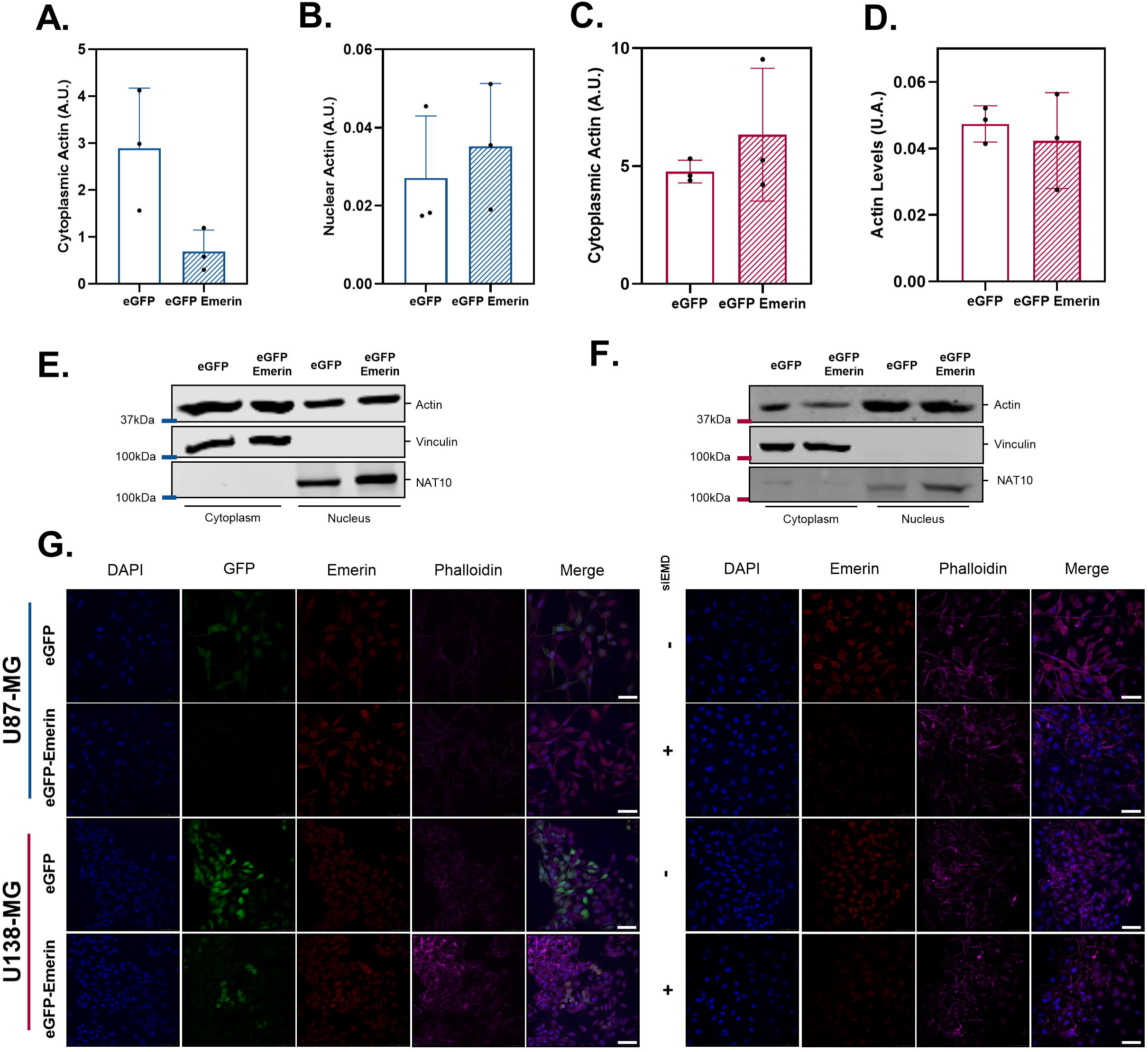
Evaluation of subcellular actin distribution following Emerin overexpression and cellular morphology after Emerin modulation in glioblastoma cells. **A–F**. Cellular fractionation of U87-MG (A-B, E) and U138-MG (C-D, F) cells transfected with eGFP or eGFP-Emerin. (A-D) Quantification of cytoplasmic and nuclear actin levels (A.U.). Data are presented as mean ± standard deviation. (E-F) Representative western blot analysis of cytoplasmic and nuclear fractions showing Actin, Vinculin (cytoplasmic marker), and NAT10 (nuclear marker) levels. **G**. Immunofluorescence analysis of cellular morphology following Emerin silencing and EGFP-Emerin expression. Cells were stained with DAPI (blue), anti-Emerin (red), and Phalloidin (magenta). GFP signals are shown in green. Representative images demonstrate alterations in cytoskeletal organization and cellular distribution following Emerin modulation. Representative micrographs were obtained upon magnification of 40x objective lenses. Scale bar = 50 µm.

### DNA damage response and repair protein levels are affected by Emerin overexpression

U87-MG and U138-MG cells overexpressing or not eGFP-EMD were treated by 10µM cisplatin and analyzed as to the expression of DDR and repair sensor proteins by 6h, 24, or 48h after. The kinetics of total p53, p-p53(Ser15) and p21^Kip1^ were discretely affected in U87-MG cells overexpressing Emerin compared to parental cells, but not for pH2AX(Ser139) what suggests defects in DNA repair mechanisms but not in DDR (Fig. 6A, 6C, 6D, 6E and 6F). Therefore, no relevant changes were observed in U138-MG cells and independently of cisplatin treatments, except by the levels of pH2AX(Ser139) that interestingly were inversely changed in Emerin overexpressing cells compared to control and after cisplatin (Fig. 6B, 6G, 6H and 6I).

**Figure 6.**
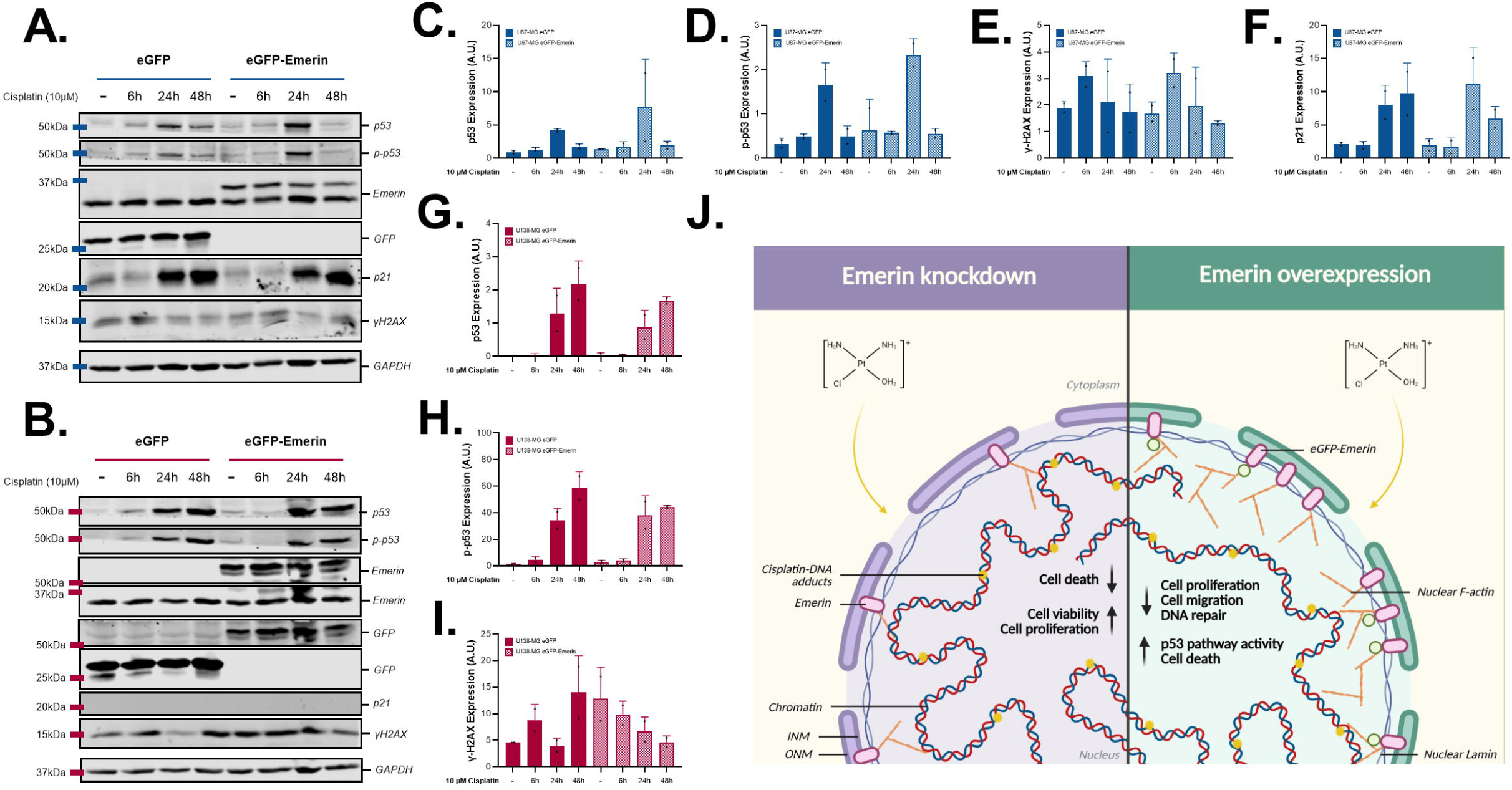
Effect of Emerin overexpression on DNA damage response and p53 pathway activation following cisplatin treatment in glioblastoma cells. A–B. Representative Western blot analysis of U87-MG (A) and U138-MG (B) cells stably expressing eGFP (control) or eGFP-Emerin, treated with 10 µM cisplatin for 6, 24, and 48 hours. Protein levels of phospho-p53 (p-p53-Ser15), total p53, p21, and pH2AX (γH2AX-Ser139) were analyzed. Emerin and GFP expression confirm stable overexpression. GAPDH was used as a loading control. **C–I** Analysis of p53 (C-D), p-p53 (E-F), pH2AX (G-H) and p21 (I) protein levels in U87-MG and U138-MG cells. Protein levels were quantified using Empiria Studio software (LI-COR), normalized to GAPDH, and expressed in arbitrary units (A.U.). Data are presented as mean ± SD. **J.** Illustrative figure representing the molecular and cellular aspects affected by Emerin modulation under conditions of cisplatin-induced genotoxic stress.

## Discussion

In this study, we demonstrated that modulation of Emerin levels directly impacts viability, survival, migration, and DNA damage response in glioblastoma cells in a p53-dependent manner. Knocking down Emerin gene expression led to increased resistance to cisplatin in U87-MG cells (wild-type p53), whereas its ectopic overexpression resulted in higher treatment sensitivity, evidenced by decreased viability, clonogenic potential, and migration. In contrast, these effects were not observed in U138-MG cells (mutant p53), indicating that Emerin’s function in this context may be strongly dependent on the functional integrity of the p53 pathway.

Recognition and repair of DNA damage are associated with dynamic changes in nuclear organization, allowing damaged regions to reposition to specific nuclear domains, facilitating recruitment of repair proteins, and influencing the choice of repair pathway [22]. These processes depend on the integration of the LINC complex, molecular motors, and nucleoskeletal components such as filamentous actin (F-actin) [23][24], which contribute to chromatin mobility and nuclear reorganization [22][25]. Nuclear F-actin formation is stimulated in response to DNA damage [23], and inhibition of its polymerization impairs repair efficiency [24]. Additionally, Emerin can bind nuclear F-actin, acting as a “pointed-end capping” protein, promoting filament stabilization and polymerization [12]. Our results demonstrate that perfect levels of Emerin expression might maintain the nuclear architecture homeostasis that favors actin polymerization/depolymerization rates ideal for cellular responses to damage. Reduced Emerin levels in glioblastoma cells are associated with greater resistance to cisplatin-induced genotoxic stress, while elevated Emerin levels promote increased treatment sensitivity. In this scenario, higher Emerin levels, by favoring F-actin stabilization may contribute to a more rigid and branched actin network that affects a controlled nuclear organization, limiting the nuclear dynamics necessary to cope with genotoxic stress, whereas its reduction may promote more resistant phenotypes.

Actin dynamics also plays a central role in regulating p53 localization and activity in response to DNA damage [26][27]. In the cytoplasm, increased actin polymerization (F-actin levels) can promote cytoplasmic retention of p53, reducing its nuclear translocation, stability, and transcriptional activity, including p21 expression [26]. Moreover, the monomeric form of actin (G-actin) is directly involved in nucleocytoplasmic transport of p53, facilitating its translocation to the nucleus, whereas the polymerized form (F-actin) also contributes to this process [27]. In this context, Emerin, by stabilizing F-actin filaments, can influence the balance between G-actin and F-actin, indirectly modulating nuclear p53 availability. This regulation is particularly relevant when considering the distinct patterns observed in the studied cell lines. In U87-MG cells, which have functional p53, actin dynamics mediated by Emerin may directly impact p53 localization and activity, modulating the DNA damage response. In contrast, U138-MG cells harbor the R175H mutation in the DNA-binding domain, which compromises p53’s transcriptional function. Thus, even if actin dynamics influence its subcellular localization, the inability to activate target genes limits the functional impact of this axis. Therefore, Emerin’s role in modulating the response to genotoxic stress may depend on functional p53 integrity, contributing to the differences observed between these two cell lines.

These findings suggest that Emerin may act as a modulator of the DNA damage response, possibly through its interaction with nuclear actin and the LINC complex, which may be influencing chromatin accessibility and repair efficiency. In this sense, alterations in levels of damage response proteins, such as p53, p-p53(Ser15), p21, and γH2AX, observed in cells overexpressing Emerin, support the hypothesis that this protein participates in fine-tuning damage signaling and repair processes.

Regarding cellular response to damage and cytoskeleton-dependent motility, the reduced migration capacity observed in cells overexpressing Emerin also aligns with studies linking nuclear envelope proteins to regulation of tumor cell motility and invasiveness [6][28]. Reduced Emerin expression is associated with smaller nuclear size and increased deformability, favoring more invasive and adaptive phenotypes [28]. Thus, elevated Emerin levels likely increase nuclear rigidity, hinder migration, particularly under chemotherapeutic stress. However, from our studies, Emerin expression was shown to be high either in wild-type or mutated p53 cells, despite their nuclear morphology appearing very different. U87-MG cells have well-defined big round oval nucleus with Emerin well distributed all over the nuclear membrane, while U183-MG cells have nucleus with a bean-like morphology, smaller and presenting some type of nuclear membrane with inner invaginations also covered by Emerin.

Additionally, these findings may relate to cytoskeletal dynamics regulation mediated by Rho family GTPases, recognized as key determinants of cell migration [17][29]. In this process, intracellular RhoA distribution changes, with higher activation in rear regions associated with contractile force generation, and evidence suggests that Emerin acts as a regulator of this process by stabilizing RhoA activation and promoting cell polarity even working on actin nucleoskeleton [29]. Differences on cytosolic actin cytoskeleton through stress fiber staining’s have also shown that Emerin knocking down seems to affect that abovementioned remodeling dynamics.

Furthermore, Emerin modulation affected GBM cell survival after cisplatin treatment. Emerin can regulate signaling pathways associated with cell proliferation by directly interacting with β-catenin and inhibiting its nuclear localization, reducing expression of Wnt/β-catenin target genes such as c-Myc and cyclin D1 [30]. Thus, elevated Emerin levels are associated with suppressed proliferation, whereas its reduction may favor proliferative programs [30], promoting genotoxic stress resistance and clonogenic potential.

Taken together, our data suggests that Emerin may act as a central regulator integrating nuclear architecture, cytoskeletal dynamics, and response to genotoxic stress through its interaction with nuclear actin and the LINC complex and may potentially serve as a biomarker of tumor aggressiveness and therapeutic sensitivity. However, the exact molecular mechanisms by which Emerin regulates specific repair pathways remain to be elucidated. Future studies should further explore Emerin’s participation in specific DNA repair pathways and its relevance for *in vivo* models. Overall, our results indicate that Emerin plays a significant role in regulating the DNA damage response and acts as a potential modulator of chemosensitivity in glioblastoma, particularly in tumors with functional p53, possibly integrating signals between the cytoskeleton and the nucleus.

## Supporting information

This supplemental file includes five supplementary figures of results that complement the results presented in the main figures.

## Acknowledgments

The authors thank Prof. Alexandre B. Cardoso (IQ-USP) for allowing the use of the Leica DMi1 and DMi8 microscopes and the technician Lilian C. Russo for help with the Leica TCS SP8 STED confocal microscope. Authors also thank technical assistance from Benedita de Oliveira and Marcia Aparecida da Silva. DJFH also thanks Yuli T. Magalhaes and Lucas F.S. Soubihe for experimental and bioinformatics support, respectively.

## Data availability statement

Data available on request from the authors.

## Contributions

Conception and design: DJFH, FLF. Development of methodology: DJFH, FLF. Acquisition of data: DJFH, FLF. Analysis and interpretation of data: DJFH, FLF. Writing, review, and/or revision of the manuscript: DJFH, FLF. Administrative, technical, or material support: FLF. Funding acquisition: FLF. Study supervision: FLF.

