## Supplementary material for "Emerin modulation impacts viability, proliferation, migration, and DNA repair signaling in cisplatin-treated glioblastoma cells": This supplemental file includes five supplementary figures of results that complement the results presented in the main figures.

**Supplementary Figures with Legends**

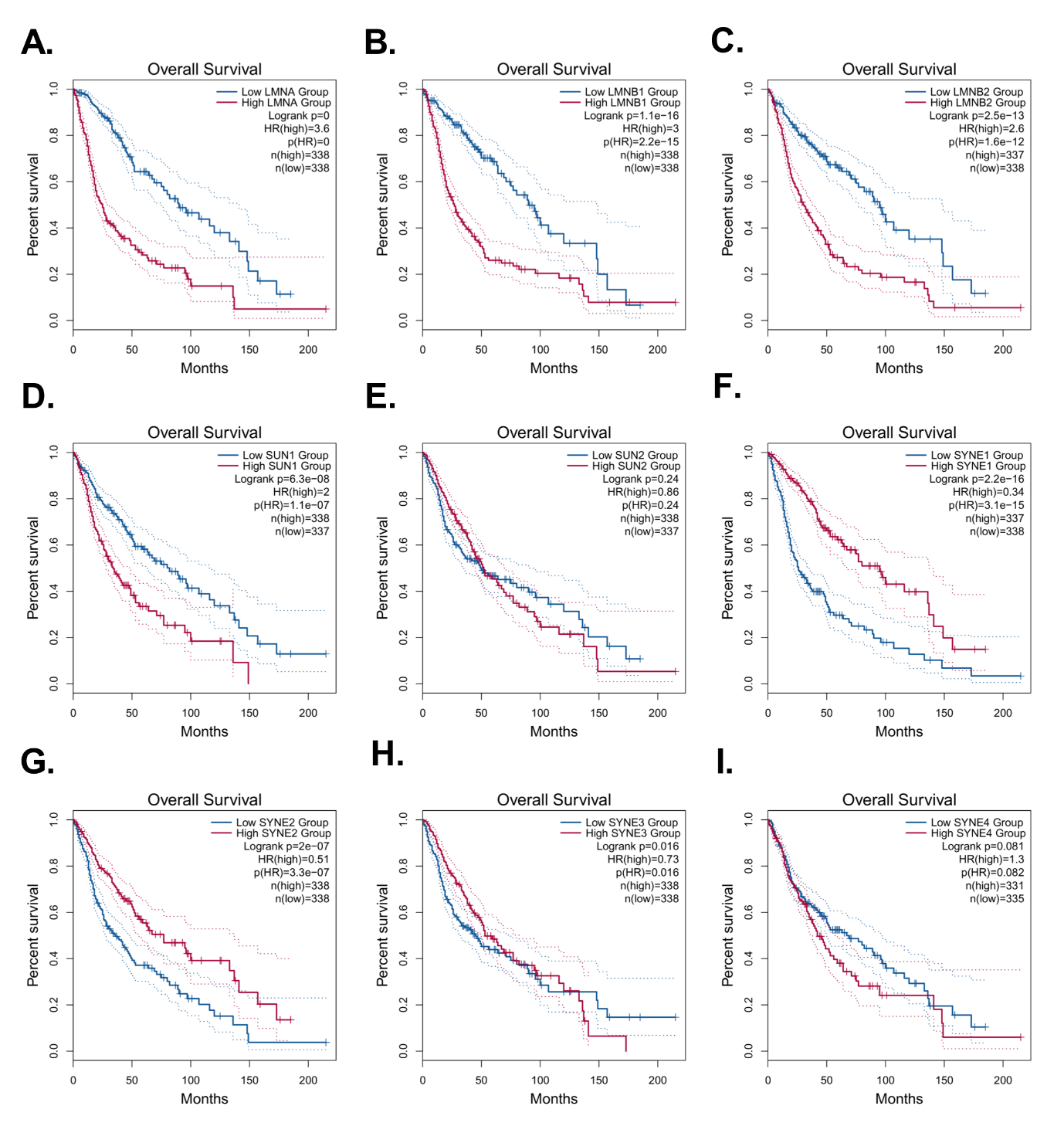

**Figure S1. Overall survival analysis of GBM and LGG patients according to the expression of LINC complex and nuclear lamina genes.** Each Kaplan–Meier plot compares high expression (red) and low expression (blue) groups for *LMNA* (A), *LMNB1* (B), *LMNB2* (C), *SUN1* (D), *SUN2* (E), *SYNE1* (F), *SYNE2* (G), *SYNE3* (H), and *SYNE4* (I). Hazard ratio (HR), log-rank p-value, and the number of patients in each group (n) are indicated in each graph. Dashed lines represent 95% confidence intervals. *LMNA: LAMIN A/C LMNB1-2: LAMIN B1-B2: SYNE1-4: NESPRIN 1-4*.

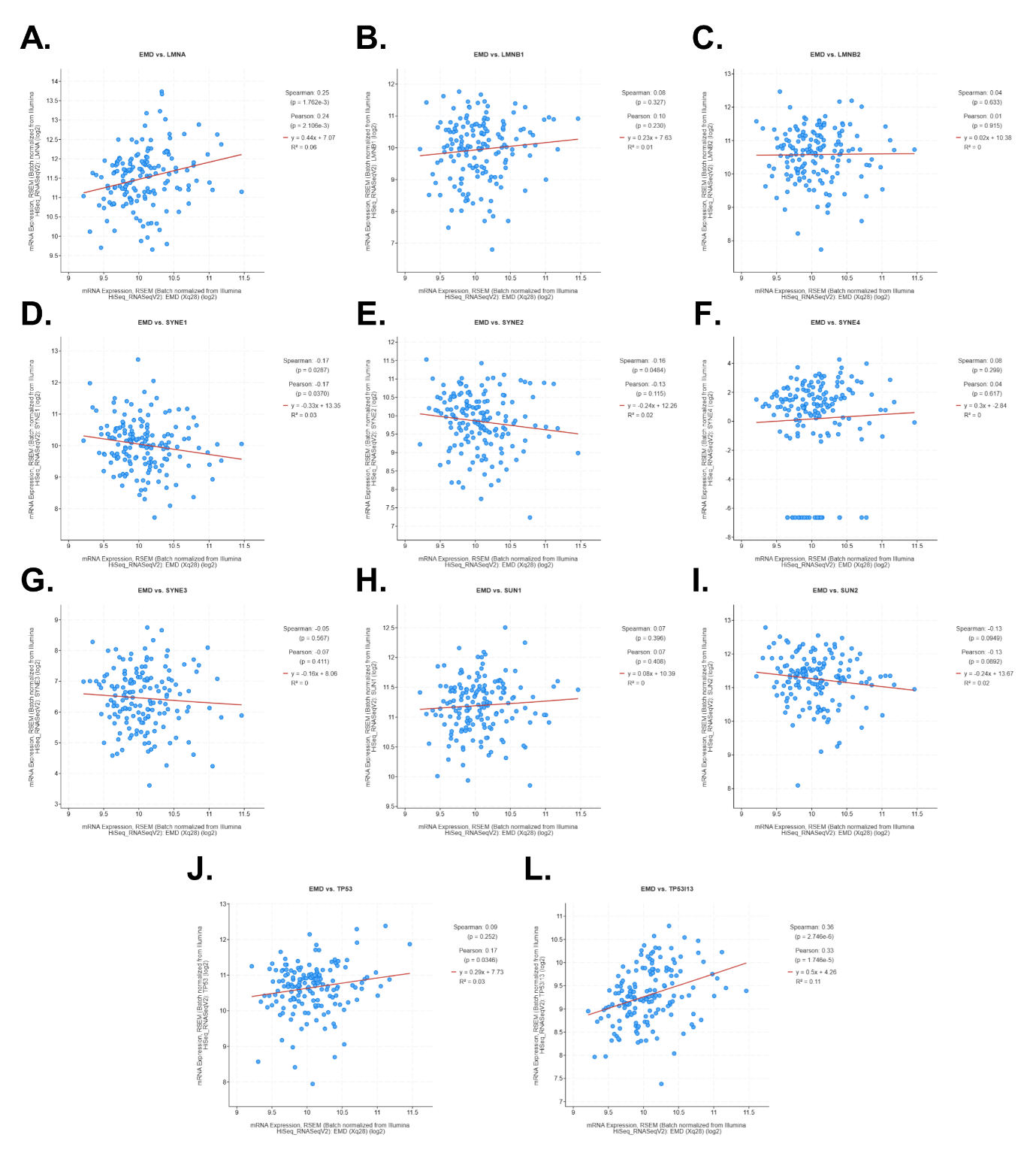

**Figure S2. Correlation between *EMERIN* (*EMD*) expression and genes associated with the nuclear lamina, cytoskeleton, and cellular response in glioblastoma samples. A–L.** Correlation analyses between EMD expression levels and the genes *LMNA* (A), *LMNB1* (B), *LMNB2* (C), *SYNE1* (D), *SYNE2* (E), *SUN1* (F), *SUN2* (G), *TP53* (J), and *TP53I13* (L) in human glioblastoma samples. Graphs show Spearman and Pearson correlation analyses, including linear regression lines and corresponding p values and correlation coefficients (r and ρ). Each dot represents an individual tumor sample.

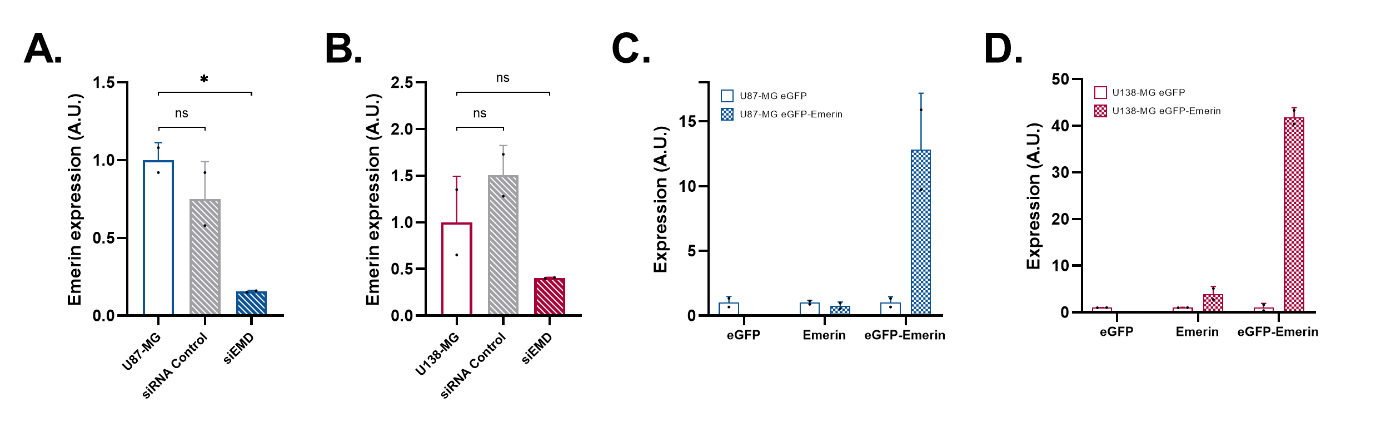

**Figure S3. Quantification of Emerin knockdown and overexpression in glioblastoma cells. A–B.** Analysis of Emerin protein levels following siRNA-mediated knockdown in U87-MG (A) and U138-MG (B) cells. Cells were transfected with control siRNA (siRNA Control) or Emerin-specific siRNA (siEMD).**C–D** Analysis of Emerin, eGFP and eGFP-Emerin protein levels in U87-MG (C) and U138-MG (D) cells stably expressing eGFP (control) or eGFP-Emerin. Protein bands were quantified using Empiria Studio software (LI-COR), normalized to GAPDH, and expressed in arbitrary units (A.U.). Data are presented as mean ± SD. Statistical significance is indicated as follows: *p < 0.05; ns, not significant.

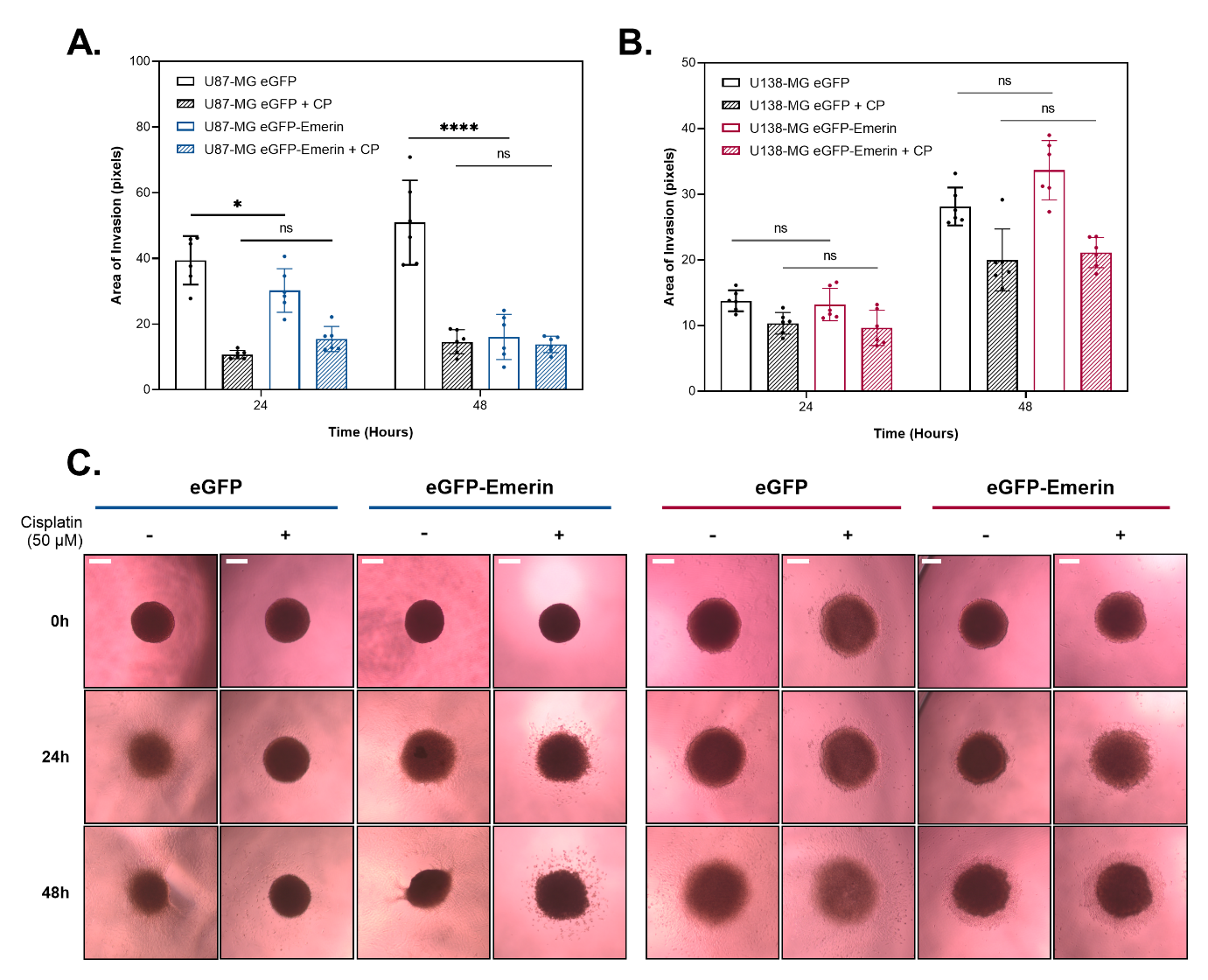

**Figure S4. Emerin modulation alters clonogenic survival following cisplatin treatment in p53 wild-type GBM cells. A–B.** Representative clonogenic assays of U87-MG (A) and U138-MG (B) cells transfected with Emerin-specific siRNA (siEMD), treated or not with cisplatin. **C–D.** Representative clonogenic assays of U87-MG (C) and U138-MG (D) cells stably expressing eGFP (control) or eGFP-Emerin, treated or not with cisplatin. Colony plates are representative of three independent experiments. Statistical analysis was performed using two-way ANOVA.

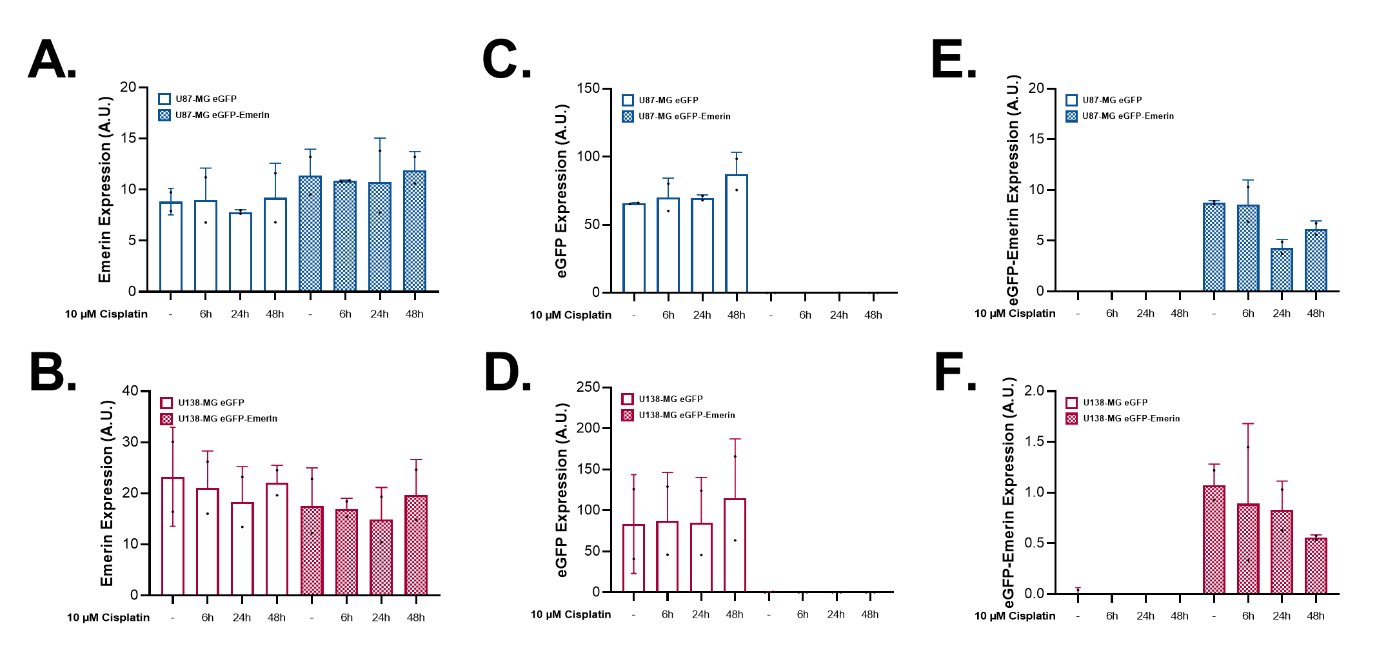

**Figure S5. Quantification of Emerin, eGFP, and eGFP-Emerin expression following cisplatin treatment.** Analysis of Emerin (A-B), eGFP (C-D) and eGFP-Emerin (E-F) protein levels in U87-MG and U138-MG cells**.** Protein levels were quantified using Empiria Studio software (LI-COR), normalized to GAPDH, and expressed in arbitrary units (A.U.). Data is presented as mean ± SD.
